# Nerve pathology is prevented by linker proteins in mouse models for *LAMA2*-related muscular dystrophy

**DOI:** 10.1101/2022.05.19.492629

**Authors:** Judith R. Reinhard, Emanuela Porrello, Shuo Lin, Pawel Pelczar, Stefano C. Previtali, Markus A. Rüegg

## Abstract

*LAMA2*-related muscular dystrophy (LAMA2 MD or MDC1A) is caused by mutations in the *LAMA2* gene encoding laminin-α2, the long chain of several heterotrimeric laminins. Laminins are essential components of the extracellular matrix that interface with underlying cells. The pathology of LAMA2 MD patients is dominated by the severe muscular dystrophy but also involves other tissues. In the *dy*^*W*^/*dy*^*W*^ mouse model for LAMA2 MD, amelioration of muscle function by skeletal muscle-specific expression of the two linker proteins, mini-agrin and αLNNd, is sufficient to greatly increase survival. In such survivors, the phenotype is dominated by the hindlimb paralysis. We now show that ubiquitous expression of the two linker proteins in *dy*^*W*^/*dy*^*W*^ mice improves muscle function and prevents hindlimb paralysis. The same ameliorating effect of the linker proteins was seen in *dy*^*3K*^/*dy*^*3K*^ mice, which represent the most severe mouse model of LAMA2 MD. In summary, these data show that the two linker proteins can compensate the loss of laminin-α2 in many, if not all tissues affected in LAMA2 MD.

## INTRODUCTION

Skeletal muscle is required to maintain posture, for locomotion, respiration and overall metabolic homeostasis. Motor neurons originating in the spinal cord control the contraction of muscle fibers by innervating them at the neuromuscular junction (NMJ) where they release acetylcholine to trigger an action potential in the muscle fiber to cause its contraction. At the same time, the length and position of muscle fibers is controlled by the proprioceptive system, which feeds this information back to the spinal cord. Thus, skeletal muscle function requires a complex network that sends impulses from the spinal cord to muscle and back again *via* peripheral nerves (Proske & Gandevia, 2012). To allow fast conductance of the action potentials, motor and proprioceptive axons are myelinated by Schwann cells.

Congenital muscular dystrophies are characterized by early onset (diagnosed around birth), high severity and reduced lifespan. *LAMA2*-related muscular dystrophy (LAMA2 MD or MDC1A) is caused by mutations in *LAMA2*, the gene coding for the laminin-α2 chain of the heterotrimeric laminins, composed of α, β and γ chains (Domogatskaya *et al*, 2012; Yurchenco, 2011). In total, 16 different laminins (i.e. heterotrimers with different chain composition) have been described in mammals. The most abundant, α2-laminin-containing heterotrimer is laminin-211 (formerly called merosin), composed of the α2, the β1 and the γ1 chains. Laminin-211 is mainly detected in the endomysial basement membrane (BM) of mature skeletal muscle fibers, in the heart and in the endoneurial BM of the peripheral nerve (Sasaki *et al*, 2002). Consistent with this expression pattern, the disease phenotype of LAMA2 MD patients is dominated by the muscle dystrophy with some involvement of the heart, the peripheral nerve and the central nervous system (Nguyen *et al*, 2019; Sarkozy *et al*, 2020). Skeletal muscle from LAMA2 MD patients and mouse models “compensate” the loss of laminin-α2 by expression of the embryonic laminin-α4 chain (Patton *et al*, 1999; Reinhard *et al*, 2017). Laminin-α4-containing laminin-411 does not bind to the *bona fide* cell surface receptors of laminin-211 and is not able to self-polymerize (Reinhard *et al*., 2017; Yurchenco *et al*, 2018) and thus is not able to functionally compensate. In an attempt to “graft” both functions of laminin-α2 onto laminin-411, two linker proteins have been designed that consist of domains of agrin (mini-agrin; mag) or laminin-α1 and nidogen-1 (αLNNd). Both proteins bind to laminin-411; mag, in addition, binds to α-dystroglycan and αLNNd induces laminin-411 polymerization (McKee *et al*, 2007; Moll *et al*, 2001). Muscle-specific, transgenic expression of either mag or αLNNd in different LAMA2 MD mouse models [i.e., *dy*^*W*^/*dy*^*W*^, *dy*^*3K*^/*dy*^*3K*^ or *dy*^*2J*^/*dy*^*2J*^ mice (Gawlik & Durbeej, 2020)], ameliorates the disease (Bentzinger *et al*, 2005; McKee *et al*, 2017; Moll *et al*., 2001). Importantly, transgenic expression of both linker proteins in skeletal muscle fibers of *dy*^*W*^/*dy*^*W*^ mice provides additive benefit, leading to a drastic prolongation of the median lifespan from less than 4 months to more than 18 months (Reinhard *et al*., 2017). However, prolongation of survival accentuates the progressive peripheral neuropathy caused by laminin-211 deficiency in Schwann cells (Reinhard *et al*., 2017), similar to what was observed by muscle-specific, transgenic expression of *Lama2* in *dy*^*W*^/*dy*^*W*^ mice (Kuang *et al*, 1998b).

In the current work, we tested whether the linker proteins could ameliorate the muscular dystrophy and prevent hindlimb paralysis when expressed ubiquitously. To this end, we generated novel transgenic mice for the two linker proteins and evaluated their disease ameliorative effect in *dy*^*W*^/*dy*^*W*^ and *dy*^*3K*^/*dy*^*3K*^ mice. We find that ubiquitous expression of the two linker proteins restores both muscle and peripheral nerve function resulting in near-normal body weight, gait and muscle force. These data thus provide unequivocal evidence that linker proteins can functionally replace laminin-211 in muscle and in the peripheral nerve.

## RESULTS

### Sciatic nerve of LAMA2 MD mice contains laminin-α4 and linker proteins localize to the endoneurial BM when expressed ubiquitously

LAMA2 MD mouse models and LAMA2 MD patients express the compensatory laminin-411 in the muscle BM (Patton *et al*., 1999; Reinhard *et al*., 2017). As the ameliorative function of mag and αLNNd in *dy*^*W*^/*dy*^*W*^ mice requires their binding to laminin-411, we stained sciatic nerve cross-sections with antibodies to laminin-α2 and laminin-α4 (Fig 1A). The endoneurial, but not the perineurial BM was strongly positive for laminin-α2 in control mice. In contrast, laminin-α4 staining was weak in the endoneurial but strong in the perineurial BM. In *dy*^*W*^/*dy*^*W*^ mice, laminin-α4 staining was strong in both endoneurial and perineurial BM (Fig 1A).

**Figure 1.**
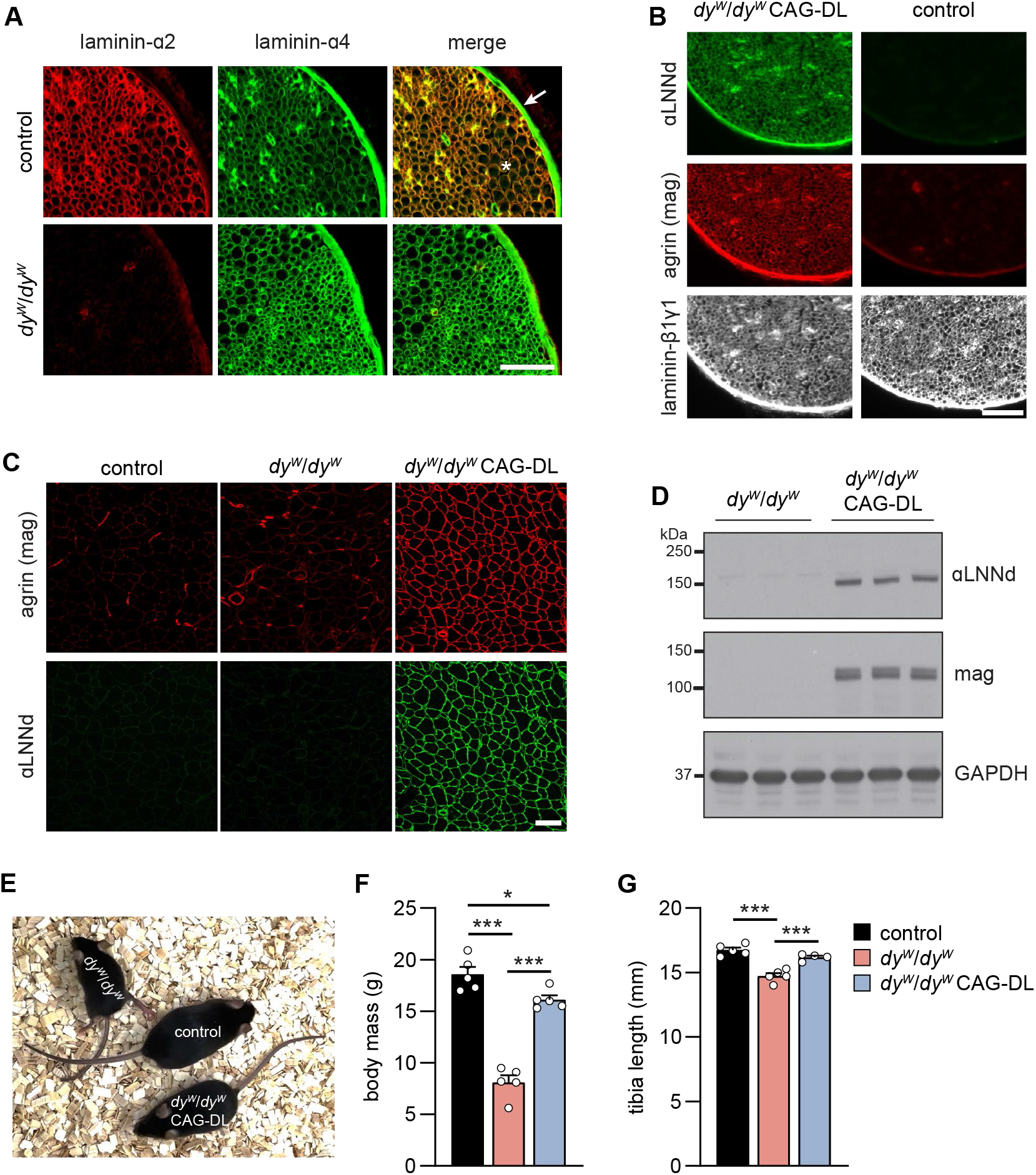
Expression of laminins and linkers in control and LAMA2 MD mice. **A** Cross-sections of the sciatic nerve from control and *dy*^*W*^/*dy*^*W*^ mice stained with antibodies to laminin-α2 and laminin-α4. Note the increase in laminin-α4 immunoreactivity in the endoneurium in *dy*^*W*^/*dy*^*W*^ mice. Arrow indicates perineurium and asterisk endoneurium. **B** – **C** Cross-sections of the sciatic nerve (**B**) and TA muscle (**C**) of 8-week-old mice of the indicated genotype. The linker proteins αLNNd and mag are present in the endoneurium and perineurium of the sciatic nerve (**B**) and in the muscle fiber basement membrane (**C**). **D** Western blot analysis to detect αLNNd and mag in lysates from *triceps brachii* muscle of 8-week-old mice. GAPDH was used as loading control. **E** Overall phenotype of 14-week-old mice of the indicated genotype. **F, G** Body mass (**F**) and tibia length (**G**) of 8-week-old mice of the indicated genotype. Data are mean ± SEM. \**P* < 0.05; \*\*\**P* < 0.001; by one-way ANOVA with Bonferroni post hoc test. Scale bars: 50 μm (**A-B**), 100 μm (**C**). N = 4-5 female mice per group.

To explore the possibility that expression of the two linker proteins mag and αLNNd would also ameliorate non-muscle phenotypes in *dy*^*W*^/*dy*^*W*^ mice, we inserted the sequences coding for mouse mag or αLNNd into the *Rosa26* locus of C57BL/6J mice by CRISPR/Cas9-driven homology-directed repair (for a schematic presentation see Fig S1A). In these knock-in mice, upon Cre-mediated deletion of the stop cassette, the linker proteins are expressed by the CAG promoter. To test the functionality of the coding sequence, we transfected COS7 cells with the targeting vectors. Only upon co-transfection with a Cre-expressing plasmid, we detected mag or αLNNd in cell extracts (top rows) and the culture medium (bottom rows) with the expected size (Fig S1B). This confirms the functionality of both constructs and shows that the loxP-Stop-loxP (LSL) cassette prevents any leakage. To select the appropriate Cre system, we crossed different mouse lines with tdTomato reporter mice [Rosa26-CAG-LSL-tdTomato (Madisen *et al*, 2010)]. One of those were the transgenic CAG-Cre-ER™ mice that express a tamoxifen-responsive Cre-recombinase under the CAG promoter (Hayashi & McMahon, 2002). Examination of skeletal muscles showed that CAG-Cre-ER™ caused strong Tdtomato expression (observed as red coloring of the entire muscle), irrespective of tamoxifen injection (Fig S1C). These experiments show that some recombination of the LSL cassette occurred in CAG-Cre-ER™-positive mice even in the absence of tamoxifen, as previously reported (Hayashi & McMahon, 2002), eventually sufficient for linker protein expression. Indeed, triple transgenic mice, not injected with tamoxifen, that were positive for the CAG-Cre-ER™ transgene and carried one copy of the Rosa26-CAG-LSL-mag and one copy of Rosa26-CAG-LSL-αLNNd allele (called CAG-DL for <u>CAG</u>-driven <u>D</u>ouble <u>L</u>inker) expressed both linker proteins from birth onwards with some difference in the expression levels between individual mice in the first two weeks (Fig EV1D). The deletion of the LSL cassette in Cre-positive mice in brain and skeletal muscle was also confirmed on the genomic level (Fig S1E). By further intercrossing, we generated *dy*^*W*^/*dy*^*W*^ CAG-DL mice and compared them with *dy*^*W*^/*dy*^*W*^ and control mice. None of the mice were injected with tamoxifen.

In *dy*^*W*^/*dy*^*W*^ CAG-DL mice, the endoneurial and the perineurial BM was strongly positive for both linker proteins (Fig 1B) as was the muscle BM (Fig 1C), indicating that the linker proteins are expressed and incorporated into the BM. Western blot analysis of muscle lysates from 8-week-old mice confirmed expression of the linker proteins in *dy*^*W*^/*dy*^*W*^ CAG-DL mice (Fig 1D). The expression of the linker proteins induced a striking overall phenotypic improvement in *dy*^*W*^/*dy*^*W*^ mice, to the extent that it became difficult to distinguish controls from *dy*^*W*^/*dy*^*W*^ CAG-DL mice (Fig 1E). Presence of the linkers caused *dy*^*W*^/*dy*^*W*^ to reach a close to normal body mass (Fig 1F) and size (Fig 1G).

### Linker proteins prevent hindlimb paralysis by restoring axonal sorting and myelination

We also noticed that in contrast to *dy*^*W*^/*dy*^*W*^ mice and *dy*^*W*^/*dy*^*W*^ mice that express the linker proteins only in skeletal muscle fibers (Reinhard *et al*., 2017), *dy*^*W*^/*dy*^*W*^ CAG-DL mice did not display paralysis of the hindlimbs (Movie S1). The peripheral neuropathy in LAMA2 MD mouse models is caused by severe defects in axonal sorting and myelination of motor and sensory axons (Feltri *et al*, 2016; Previtali & Zambon, 2020). Staining of semithin cross-sections of the sciatic nerve with toluidine blue revealed many bundles or islands of “naked” axons in *dy*^*W*^/*dy*^*W*^ mice (white arrows in Fig 2A and black arrows in Fig S2A). These large regions of non-myelinated axons were also a striking feature in the sciatic nerve of *dy*^*W*^/*dy*^*W*^ mice in ultrathin, electron microscopic (EM) pictures (Fig 2A). While these bundles were usually large in *dy*^*W*^/*dy*^*W*^ mice, there were only few of them in *dy*^*W*^/*dy*^*W*^ CAG-DL mice and they were smaller (Fig 2A; Fig S2A). Axons of *dy*^*W*^/*dy*^*W*^ mice in the bundles were naked and not wrapped by Schwann cells (white arrowheads in *dy*^*W*^/*dy*^*W*^ panel of Fig 2A) and they contained many large-caliber axons (white asterisks in *dy*^*W*^/*dy*^*W*^ panel of Fig 2A). In contrast, bundles of “naked” axons in *dy*^*W*^/*dy*^*W*^ CAG-DL mice were reminiscent of the Remak bundles in controls, which are small axons that are ensheathed by non-myelin-forming Schwann cells (blue arrowheads in control and *dy*^*W*^/*dy*^*W*^ CAG-DL panels of Fig 2A). In *dy*^*W*^/*dy*^*W*^ mice, quantification demonstrated a significant increase in the area of the bundles (Fig 2B), in the number of axons within such bundles (Fig 2C), in the number of large-caliber axons (Fig 2D) and a significant decrease in the relative number of axons fully wrapped by Schwann cells (Fig 2E) compared to controls. Importantly, all these parameters were restored to control values in *dy*^*W*^/*dy*^*W*^ CAG-DL mice. We also observed changes in the size distribution of axons in *dy*^*W*^/*dy*^*W*^ mice as compared to controls, and some improvements in the *dy*^*W*^/*dy*^*W*^ CAG-DL mice (Fig S2B). The G-ratio, an index of myelination, did not significantly change between *dy*^*W*^/*dy*^*W*^ and *dy*^*W*^/*dy*^*W*^ CAG-DL mice (Fig S2C-D). Overall, these data demonstrate that expression of the two linker proteins in the peripheral nerve largely restores axonal sorting, a process that is highly perturbed in all mouse models of LAMA2 MD, irrespective of the mutation in *Lama2* (Gawlik & Durbeej, 2020).

**Figure 2.**
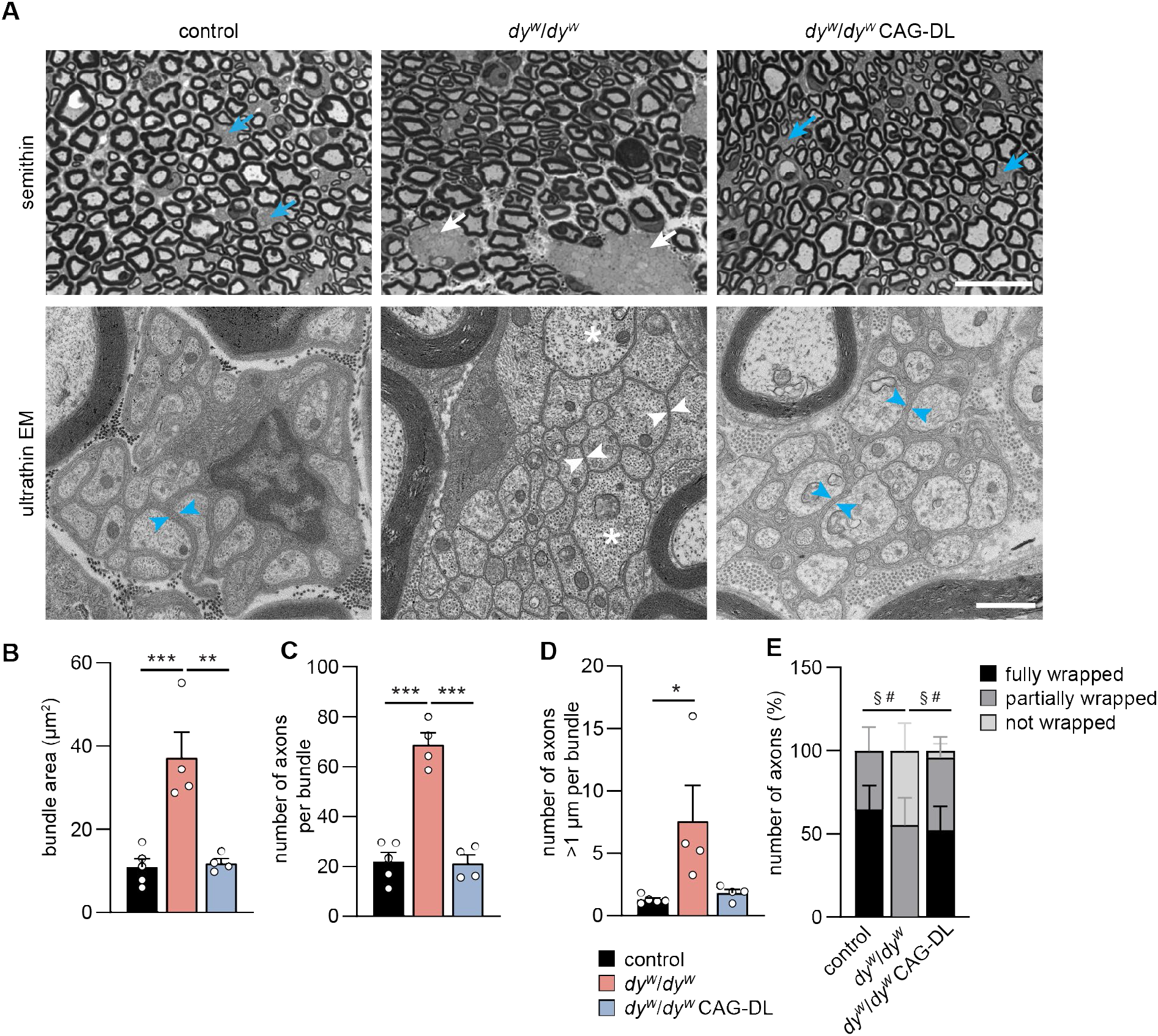
Ubiquitous expression of the linker proteins improves nerve pathology. **A** Images of semithin and ultrathin cross-sections of the sciatic nerve from 8-week-old mice of the indicated genotype. Semithin: Regions/bundles containing non-myelinated axons are indicated with arrows. In *dy*^*W*^/*dy*^*W*^ mice, the bundles are large (white arrows), while bundles are small in control and *dy*^*W*^/*dy*^*W*^ CAG-DL mice (blue arrows). Ultrathin EM: Axons within the bundles in *dy*^*W*^/*dy*^*W*^ mice are not ensheathed by Schwann cells (white arrowheads) and can be > 1 μm in diameter (asterisks). In control mice, bundles represent Remak bundles of slow-conducting axons that are ensheathed by non-myelinating Schwann cells (blue arrowheads). In *dy*^*W*^/*dy*^*W*^ CAG-DL mice, non-myelinated axons are also surrounded by non-myelinating Schwann cell processes (blue arrowheads), reminiscent of Remak bundles in control mice. **B – D** Quantitative assessment of the bundles in the different genotypes by measuring the area (**B**), the number of axons (**C**) and the number of axons > 1 μm in diameter (**D**). **E** Quantification of the extent of Schwann cell wrapping of axons in bundles in the different genotypes. Data are mean ± SEM. \**P* < 0.05; \*\**P* < 0.01; \*\*\**P* < 0.001; by one-way ANOVA with Bonferroni post hoc test (**B-D**). ^§^*P* < 0.05 in fully wrapped axons; ^#^*P* < 0.05 in not wrapped axons; by one-way ANOVA with Bonferroni post hoc test. Scale bars: 20 μm (semithin) and 1 μm (ultrathin). N = 4-5 mice per group.

### Linker proteins improve muscle function in *dy*^*W*^/*dy*^*W*^

To see whether the linker proteins improved muscle histology and function, we next examined fore- and hindlimb muscles of 8-week-old mice. Similar to the results obtained with skeletal muscle-specific expression of the two linkers (Reinhard *et al*., 2017), *triceps brachii* muscle (TRC) was strongly improved in mass, median fiber diameter and fiber size distribution (Fig S3A-C). Importantly, absolute and specific forelimb grip strength were increased such that *dy*^*W*^/*dy*^*W*^ CAG-DL developed more than 2 times more force than *dy*^*W*^/*dy*^*W*^ mice (Fig S3D). Assessment of histology in *tibialis anterior* (TA) muscle, which is affected by the hindlimb paralysis, showed strong overall improvement in H&E staining and in the extent of fibrosis, stained by Sirius Red, in *dy*^*W*^/*dy*^*W*^ CAG-DL mice compared to *dy*^*W*^/*dy*^*W*^ mice (Fig 3A). The mass of the TA almost tripled in *dy*^*W*^/*dy*^*W*^ CAG-DL mice compared to *dy*^*W*^/*dy*^*W*^ mice (Fig 3B). This increase was based on an increase in the median fiber size and in the total fiber number in *dy*^*W*^/*dy*^*W*^ CAG-DL mice, reaching close-to-normal values (Fig 3C). *Ex vivo* force measurement of the *extensor digitorum longus* (EDL) hindlimb muscle showed that absolute and specific twitch force in *dy*^*W*^/*dy*^*W*^ CAG-DL mice was improved to control levels (Fig 3D). Similarly, absolute and specific tetanic force was improved in *dy*^*W*^/*dy*^*W*^ CAG-DL mice to the extent that they were not significantly different from controls (Fig 3E). This almost complete restoration of hindlimb muscle force in *dy*^*W*^/*dy*^*W*^ CAG-DL mice is striking and is substantially better than in *dy*^*W*^/*dy*^*W*^ mice that express the linker proteins only in skeletal muscle (Reinhard *et al*., 2017), suggesting that the stronger improvement in *dy*^*W*^/*dy*^*W*^ CAG-DL mice reflects lack of atrophy in the hindlimb muscle. Together, these data show that ubiquitous expression of the two linker proteins in *dy*^*W*^/*dy*^*W*^ mice largely prevents the muscular dystrophy and the neuropathy-caused atrophy in hindlimb muscles.

**Figure 3.**
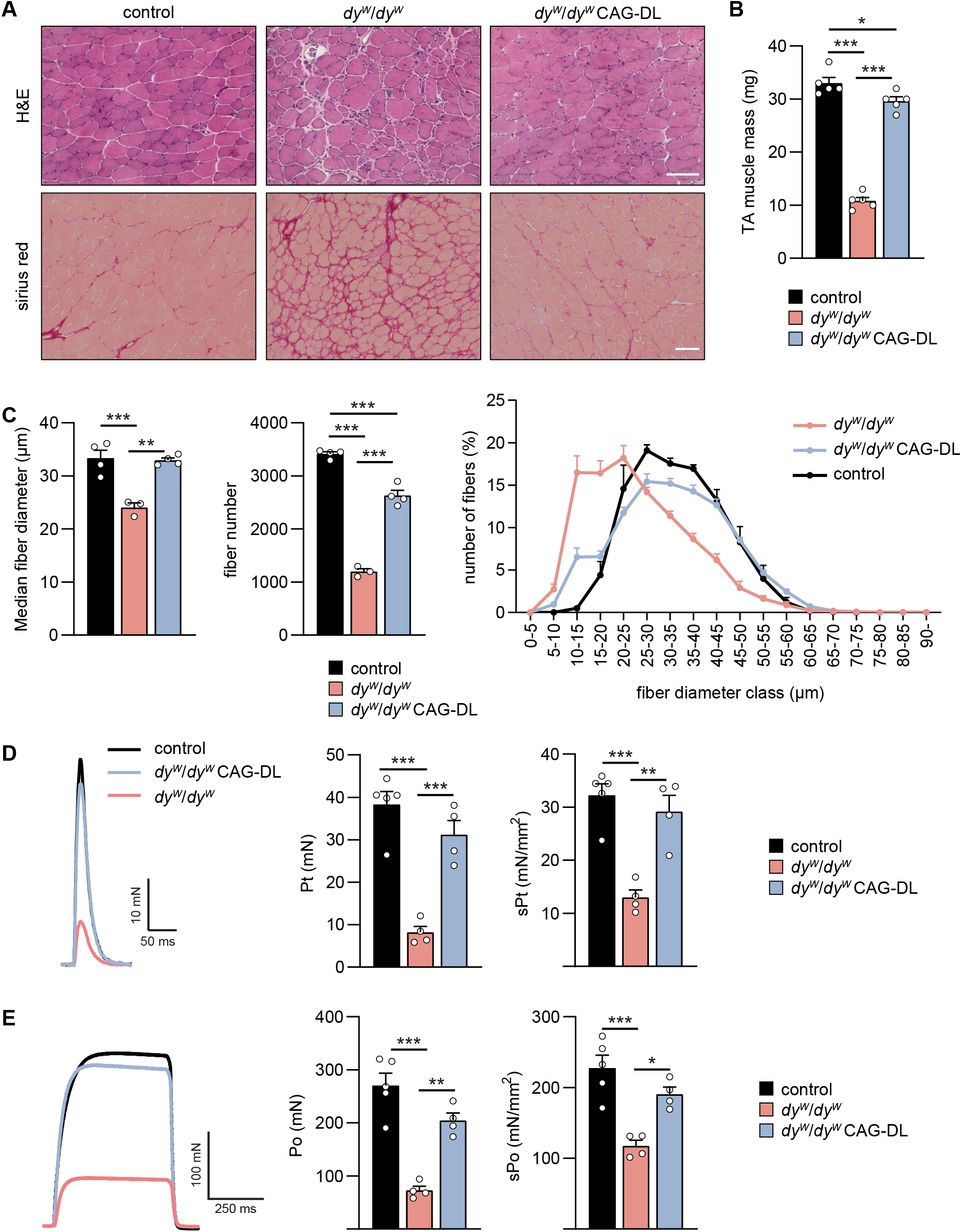
Linker proteins improve muscle histology and function in fore- and hindlimbs of *dy*^*W*^/*dy*^*W*^ mice. **A** Hematoxylin and Eosin (H&E) and sirius red staining of 8-week-old *tibialis anterior* (TA) muscle of the indicated genotypes. **B** Quantification of the TA mass from 8-week-old mice of the indicated genotypes. **C** Quantification of TA muscle fiber diameter and number from 8-week-old mice of the indicated genotypes. **D, E** Twitch (**D**) or tetanic force (**E**) of *extensor digitorum longus* (EDL) muscle from 8-week-old mice of the indicated genotypes. Shown are representative force traces (left), absolute (middle) and specific (right) force. Data are mean ± SEM. \**P* < 0.05; \*\**P* < 0.01; \*\*\**P* < 0.001; by one-way ANOVA with Bonferroni post hoc test. N = 3-5 female mice per group.

### Profound motor improvement by linker proteins is sustained in different LAMA2 MD mouse models

To assess the functional improvement in peripheral nerves in *dy*^*W*^/*dy*^*W*^ CAG-DL mice we performed quantitative assessment of locomotion and gait (Fig 4A). Average speed of *dy*^*W*^/*dy*^*W*^ mice was less than half of that of controls but significantly improved by the expression of the two linker proteins (Fig 4B). To get an estimate on the usage of front- and hindlimbs, we analyzed the weight put onto each leg by measuring the contact area of the paws (Fig 4C). In control and *dy*^*W*^/*dy*^*W*^ CAG-DL mice, front- and backpaw contact areas were of similar size, whereas they were much smaller in *dy*^*W*^/*dy*^*W*^ mice (due to the reduced body mass and size). To control for this body mass and size difference, we compared the ratio of the contact area of the back versus front paws. While the ratio was close to one in control and *dy*^*W*^/*dy*^*W*^ CAG-DL mice, it was significantly reduced in *dy*^*W*^/*dy*^*W*^ mice (Fig 4C). Thus, hindlimb paralysis prevents loading of the hindlegs in *dy*^*W*^/*dy*^*W*^ and thus reduces the relative contact area, while *dy*^*W*^/*dy*^*W*^ CAG-DL mice use all limbs similarly. Finally, motor coordination and muscle function was assessed by the rotarod assay. Eight-week-old *dy*^*W*^/*dy*^*W*^ mice showed a very short latency to fall, while control or *dy*^*W*^/*dy*^*W*^ CAG-DL mice managed to stay on the rod for a similar time (Fig 4D). Importantly, the performance of *dy*^*W*^/*dy*^*W*^ CAG-DL mice remained similar to control mice at 5 months of age (Fig 4D, right). Together, these data show that the ameliorating effect of the two linker proteins in the peripheral nerve allows for a close-to-normal locomotor function.

**Figure 4.**
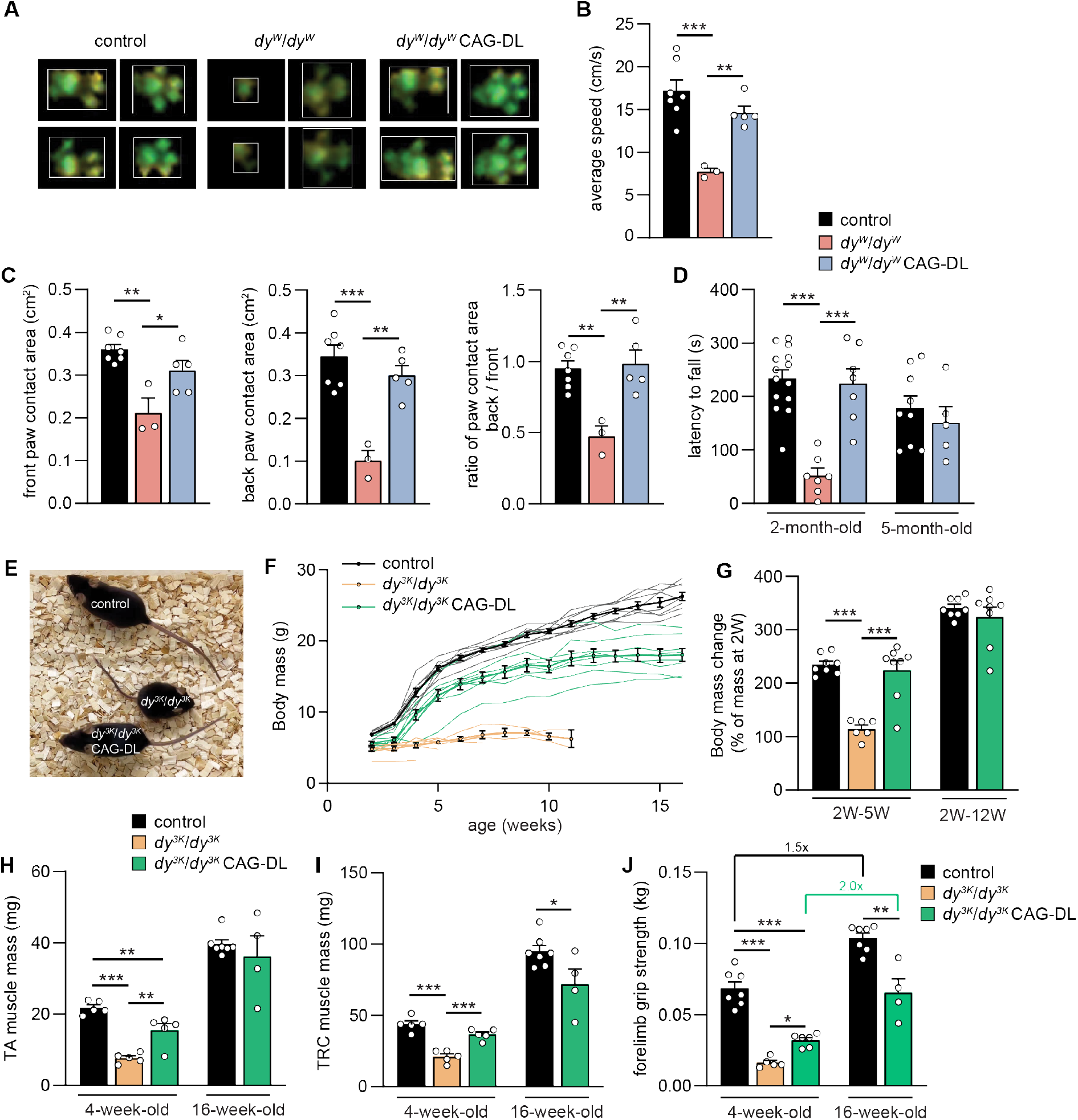
Motor performance of *dy*^*W*^/*dy*^*W*^ and evaluation of the treatment effect in *dy*^*3K*^/*dy*^*3K*^ mice. **A** Representative paw prints of mice of the indicated genotypes using a gait analysis system (CatWalk). **B, C** Quantitative assessment of locomotor speed (**B**) and paw contact area (**C**) using a gait analysis system (CatWalk). **D** Rotarod-based assessment of motor function and coordination in 2- and 5-month-old mice of the indicated genotype. Note that *dy*^*W*^/*dy*^*W*^ mice are not included at 5 months because they usually die before. **E** Overall phenotype of 4-week-old mice of the indicated genotype. **F** Body mass curves of female mice from 2 weeks to 15 weeks of age of the indicated genotype. Curves for individual mice are shown including the mean ± SEM of each cohort (thick line). Body mass curves for individual *dy*^*3K*^/*dy*^*3K*^ mice are discontinuous as all of the mice died at the age between 3 and 11 weeks. **G** Relative increase in body mass from 2 weeks of age until 5 and 12 weeks, respectively. Note that weight gain in *dy*^*3K*^/*dy*^*3K*^ CAG-DL mice up to 12 weeks is similar to controls. **H, I** Muscle mass of *tibialis anterior* (TA) and *triceps brachii* (TRC) of female mice at different age. **J** Forelimb grip strength of female mice of the indicated genotypes at 4 and 16 weeks of age. Data are mean ± SEM. \**P* < 0.05; \*\**P* < 0.01; \*\*\**P* < 0.001; by one-way ANOVA with Bonferroni post hoc test. Scale bars: 5 mm. N = 3-8 mice per group.

Next, we tested whether expression of the two linker proteins would also improve the phenotype (including the peripheral neuropathy) in the most severe mouse model for LAMA2 MD by crossing the CAG-DL triple transgene combination into *dy*^*3K*^/*dy*^*3K*^ mice (Miyagoe *et al*, 1997). Like *dy*^*W*^/*dy*^*W*^ mice, the dystrophic phenotype in *dy*^*3K*^/*dy*^*3K*^ mice develops around birth, they show a strong hindlimb paralysis, but a shorter lifespan than *dy*^*W*^/*dy*^*W*^ mice (Gawlik & Durbeej, 2020; Yurchenco *et al*., 2018). Because the median survival of the *dy*^*3K*^/*dy*^*3K*^ mice reaches only 3-5 weeks (Yurchenco *et al*., 2018), we compared all genotypes at 4 weeks of age and conducted a second analysis of controls and *dy*^*3K*^/*dy*^*3K*^ CAG-DL mice at 16 weeks of age. First, we quantified laminin-α4 expression in *dy*^*3K*^/*dy*^*3K*^ mice by Western blot analysis of TRC muscles from 4-week-old mice. Laminin-α4 was increased to a similar extent as in *dy*^*W*^/*dy*^*W*^ mice (Fig S4A), confirming prior data using immunofluorescence on muscle cross-sections (Bentzinger *et al*., 2005; Gawlik *et al*, 2004).

As in *dy*^*W*^/*dy*^*W*^ mice, expression of the linker proteins strongly improved the overall disease phenotype (Fig 4E; Movie S2). Until weaning at three weeks, body mass of *dy*^*3K*^/*dy*^*3K*^ and *dy*^*3K*^/*dy*^*3K*^ CAG-DL mice was similar but there was a clear separation between the two genotypes starting at 4 weeks (Fig 4F). While *dy*^*3K*^/*dy*^*3K*^ mice did not thrive at all from 2 to 5 weeks of age, controls and *dy*^*3K*^/*dy*^*3K*^ CAG-DL mice more than doubled their body mass (Fig 4G, left). The relative increase in body mass from 2 weeks to 12 weeks remained similar between control and *dy*^*3K*^/*dy*^*3K*^ CAG-DL mice (Fig 4G, right). These data indicate that the linker proteins do not prevent the early disease onset in *dy*^*3K*^/*dy*^*3K*^ mice until weaning but then allow the mice to thrive. In line with this, we observed some growth deficits at 4 weeks of age but a close to normal size of *dy*^*3K*^/*dy*^*3K*^ CAG-DL at the age of 16 weeks (Fig S4B). Histology of TA muscle was improved in *dy*^*3K*^/*dy*^*3K*^ CAG-DL mice compared to *dy*^*3K*^/*dy*^*3K*^ mice (Fig S4C). The number of muscle fibers as well as their size was increased in *dy*^*3K*^/*dy*^*3K*^ CAG-DL mice compared to *dy*^*3K*^/*dy*^*3K*^ mice (Fig S4D). At 4 weeks of age, mass of TA and TRC muscles from *dy*^*3K*^/*dy*^*3K*^ CAG-DL mice were significantly higher than in *dy*^*3K*^/*dy*^*3K*^ mice and muscles of linker-expressing *dy*^*3K*^/*dy*^*3K*^ mice continued to gain mass (Fig 4H, I). Functionally, grip strength was significantly higher in 4-week-old *dy*^*3K*^/*dy*^*3K*^ CAG-DL mice compared to *dy*^*3K*^/*dy*^*3K*^ mice and their grip strength increased from 4 to 16 weeks to a similar extent as in controls (Fig 4J). When grip strength was normalized to the body mass, 4-week-old *dy*^*3K*^/*dy*^*3K*^ CAG-DL mice were not significantly stronger than *dy*^*3K*^/*dy*^*3K*^ mice (Fig S4E), which might be due to the rather late onset of the improvement (see body mass curve in Fig 4F). However, the normalized forelimb grip strength of *dy*^*3K*^/*dy*^*3K*^ CAG-DL and controls were not different anymore at the age of 16 weeks (Fig S4E). Consistent with the improvement in grip strength, *ex vivo* tetanic and twitch force of EDL muscle was significantly higher in *dy*^*3K*^/*dy*^*3K*^ CAG-DL mice than in *dy*^*3K*^/*dy*^*3K*^ mice (Fig S4F, G) and further increased from 4 to 16 weeks. Finally, the overall phenotypic improvement in mice expressing the linker proteins was maintained and mice did not develop any hindlimb paralysis (Movie S3; 2-month-old and 16-month-old *dy*^*3K*^/*dy*^*3K*^ CAG-DL mouse). Taken together, these data demonstrate that the linker proteins largely restore the functionality of skeletal muscle and the peripheral nerve in the two most severe mouse models for LAMA2 MD.

## DISCUSSION

This work provides evidence that ubiquitous expression of the two linker proteins mini-agrin (mag) and αLNNd is able to correct the disease phenotype in both the skeletal muscle and the peripheral nerve of laminin-α2 deficient mice (*dy*^*W*^/*dy*^*W*^ and *dy*^*3K*^/*dy*^*3K*^). Because of the short lifespan of *dy*^*3K*^/*dy*^*3K*^ mice, we analyzed the sciatic nerves only in *dy*^*W*^/*dy*^*W*^ mice. As *dy*^*3K*^/*dy*^*3K*^ CAG-DL do not show any hindlimb paralysis (oldest mouse analyzed at 16 months), we suggest that the peripheral neuropathy is also restored in this mouse model.

Besides the severe muscular dystrophy, the peripheral neuropathy is a strong contributor to the overall phenotype in mouse models for LAMA2 MD, as it manifests as a partial paralysis of the hindlimbs. Mechanistic studies have shown that this phenotype is likely based on the failure to properly sort the axons according to their size, a process that is driven by Schwann cells and starts in rodents approximately one week before birth and finishes two weeks after birth (Feltri *et al*., 2016). We examined the peripheral nerves of 8-week-old mice for this phenotype by comparing the presence of non-sorted, naked axons in *dy*^*W*^/*dy*^*W*^ and *dy*^*W*^/*dy*^*W*^ CAG-DL mice. We find that this disease phenotype is largely prevented by the expression of the linker proteins and remains stable as indicated by the rotarod performance of *dy*^*W*^/*dy*^*W*^ CAG-DL at the age of 5 months and the overall locomotory behavior of a 16-month-old *dy*^*3K*^/*dy*^*3K*^ CAG-DL mouse. As muscle-specific transgenic expression of the linker proteins already increases median survival of *dy*^*W*^/*dy*^*W*^ mice from less than four months to more than eighteen months (Reinhard *et al*., 2017), we did not determine survival of *dy*^*W*^/*dy*^*W*^ CAG-DL or *dy*^*3K*^/*dy*^*3K*^ CAG-DL mice. However, our mouse colony includes several *dy*^*3K*^/*dy*^*3K*^ CAG-DL mice that are older than one year, indicating that the linker proteins also have a tremendous beneficial effect on survival in *dy*^*3K*^/*dy*^*3K*^ mice that have a reported median lifespan between three to five weeks (Gawlik & Durbeej, 2020; Yurchenco *et al*., 2018).

We know from previous studies that expression of both linker proteins adds benefit to skeletal muscle compared to the expression of only one of the linkers (Reinhard *et al*., 2017). In the current study we did not test whether this is also the case for the peripheral nerve. Results using adeno-associated virus (AAV)-based delivery of mini-agrin in *dy*^*W*^/*dy*^*W*^ mice and using CMV or CB promoters to drive expression, suggest that mini-agrin participates in the improvement of the nerve pathology (Qiao *et al*, 2018). On the other hand, evidence from other LAMA2 MD mouse models that express truncated forms of laminin-α2, suggest a major role laminin-211 polymerization in axonal sorting. These LAMA2 MD models all present with a severe hindlimb paralysis and a much less severe muscular dystrophy than *dy*^*W*^/*dy*^*W*^ or *dy*^*3K*^/*dy*^*3K*^ mice. These are the *dy*^*2J*^/*dy*^*2J*^ mice, which express reduced levels of an amino-terminally-truncated laminin-α2 subunit (Xu *et al*, 1994) and the dy^nfm417^/dy^nfm417^ mice, which carry a point mutation causing the conversion of an essential cysteine residue to arginine in the LN domain (required for polymerization) of laminin-α2 (Patton *et al*, 2008). Hence, the peripheral neuropathy in those mouse models is likely based on the compromised polymerization of the mutated laminin-211. If this holds true, expression of αLNNd could be the main contributor to the observed improvement of the neuropathy in *dy*^*W*^/*dy*^*W*^ CAG-DL and *dy*^*3K*^/*dy*^*3K*^ CAG-DL mice. It will be interesting to address this question in future.

The sequence coding for mini-agrin or αLNNd are small enough to be packaged into AAV vectors, which have a maximal capacity for foreign DNA of approximately 5 kb. Thus, AAV-mediated delivery of the two linker proteins might be a possible way to treat LAMA2 MD patients. Our finding that ubiquitous expression of the two linker proteins ameliorates both, the muscle and nerve phenotype suggests that such a gene therapeutic approach might be feasible using a ubiquitous promoter to drive expression of mini-agrin and/or αLNNd. However, mutations in *LAMA2* in humans manifests largely as a muscular dystrophy and there are only few patients that are reported to suffer from a peripheral neuropathy (Sarkozy *et al*., 2020). Thus, muscle-specific promoters to drive expression of the linker proteins may have the advantage of causing less off-target toxicity of the transgenes. It is of course possible that the treatment of the LAMA2 MD patients and the subsequent amelioration of the muscular dystrophy by the linker proteins may make the peripheral neuropathy more apparent. Whether it is an axonal sorting defect that underlies these occasional peripheral neuropathies in human patients remains an open question, largely due to the lack of nerve biopsies from LAMA2 MD patients. Axonal sorting defects have only recently been described in autopsy samples from infantile tissue of spinal muscular atrophy (SMA) patients (Kong *et al*, 2021). In SMA, an AAV9-based gene therapy using a ubiquitous promoter to express the missing SMN1 protein was recently approved (Hoy, 2019). Although the cells that are involved in the axonal sorting defect differ between SMA (motor neurons) and LAMA2 MD (Schwann cells), the human data from SMA patients suggest that early AAV-based intervention may be important to prevent this sorting defect. Hence, besides the treatment of the muscular dystrophy, early treatment of LAMA2 MD patients may also prevent a possible axonal sorting defect.

Recent evidence from gene therapy trails in Duchenne Muscular Dystrophy (DMD) patients indicate that the expression of micro-dystrophin may cause some immune reaction against the transgene (Manini *et al*, 2021). In contrast to the situation in DMD patients who do not express dystrophin, mini-agrin and αLNNd are assembled from domains of proteins that are expressed by LAMA2 MD patients and thus they should not cause strong immune responses. Moreover, both linker proteins do not require intracellular association with other proteins but are secreted from the cells that express them and exert their function in the extracellular matrix. Hence, they can also be incorporated into the BM adjacent to non-expressing cells. This feature may also be a substantial advantage in gene therapeutic approaches as it may allow lowering the dose needed to treat patients. In summary, the data presented here using the newly generated transgenic mice further supports the concept of AAV-based gene therapy as a viable treatment option for LAMA2 MD patients. Our data open the possibility to use ubiquitous expression of the linker proteins so that all tissues that might be affected in the disease are reached.

## Materials and Methods

### Mice

As mouse models for LAMA2 MD we used *dy*^*W*^/*dy*^*W*^ mice (Kuang *et al*, 1998a; Kuang *et al*., 1998b) [B6.129S1(Cg)-*Lama2*^*tm1Eeng*^/J; available from the Jackson Laboratory] and *dy*^*3K*^*/dy*^*3K*^ mice [B6.129P2(Cg)-*Lama2*^*tm1Stk*^, a kind gift from Drs. Shin’ichi Takeda and Yuko Miyagoe-Suzuki] (Miyagoe *et al*., 1997). Genotyping was performed as described previously for *dy*^*W*^/*dy*^*W*^ mice (Kuang *et al*., 1998b) and *dy*^*3K*^*/dy*^*3K*^ mice (Bentzinger *et al*., 2005).

Transgenic knock-in mag mice [C57BL/6J-Gt(ROSA)26Sor^tm1(CAG-mag)Rueg^] and αLNNd mice [C57BL/6J-Gt(ROSA)26Sor^tm2(CAG-aLNNd)Rueg^] were generated by CRISPR-Cas9 mediated knockin into the *Rosa26* locus. To make mag resistant to protein cleavage, we introduced the single amino acid substitution K793A into the sequence encoding the mouse version of mag (mini-agrin) (Meinen *et al*, 2007) by PCR-based site directed mutagenesis. Nucleotide sequences for mag and αLNNd (McKee *et al*., 2017) were subcloned by Gibson Assembly into the Ai9 Rosa26 targeting vector (Madisen *et al*., 2010); Addgene Cat. 22799) by replacing tdTomato. The diphtheria toxin (DTA) cassette in the Ai9 plasmid was removed by restriction digest.

Transgene integration was carried out using <u>t</u>argeted integration with linearized <u>d</u>sDNA CRISPR [Tild-CRISPR (Yao *et al*, 2018)] in fertilized mouse oocytes. Linearized dsDNA fragments used for targeting contained the transgene cassette flanked by ∼800 bp-long homology arms targeting the canonical *XbaI* site in the first intron of the *Rosa26* locus. crRNA targeting the overlapping sequence 5’-ACT CCA GTC TT<u>T CTA GA</u>A GA**T GG** (*XbaI* site underlined, PAM sequence in bold) (Chu *et al*, 2016) was used to generate cr:trcrRNA-Cas9 RNPs capable of inducing DNA double-strand breaks (DSB) and subsequent homologous recombination at the target site. C57BL/6J female mice underwent ovulation induction by intraperitoneal (i.p.) injection of 5 IU equine chorionic gonadotrophin (PMSG; Folligon–InterVet), followed by i.p. injection of 5 IU human chorionic gonadotropin (Pregnyl–Essex Chemie) 48 h later. For the recovery of zygotes, C57BL/6J females were mated with males of the same strain immediately after the administration of human chorionic gonadotropin. All zygotes were collected from oviducts 24 h after the human chorionic gonadotropin injection and were then freed from any remaining cumulus cells by a 1-2 min treatment of 0.1% hyaluronidase (Sigma-Aldrich) dissolved in M2 medium (Sigma-Aldrich). Mouse embryos were cultured in M16 (Sigma-Aldrich) medium at 37°C and 5% CO_2_. For micromanipulation, embryos were transferred into M2 medium. All microinjections were performed using a microinjection system comprised of an inverted microscope equipped with Nomarski optics (Nikon), a set of micromanipulators (Narashige), and a FemtoJet microinjection unit (Eppendorf). Injection solution containing: 100 ng/μl (60 μM) Cas9 protein (IDT), 100 μM cr:trcrRNA Rosa26 (IDT) and 20 ng/μl linearized dsDNA was microinjected into the male pronuclei of fertilized mouse oocytes until 20-30% distension of the organelle was observed. Embryos that survived the microinjection were transferred on the same day into the oviducts of 8-16-week-old pseudopregnant Crl:CD1 (ICR) females (0.5 d used after coitus) that had been mated with sterile, genetically-vasectomized males (Haueter *et al*, 2010) the day before embryo transfer. Pregnant females were allowed to deliver and raise their pups until weaning age.

Targeted pups were screened by PCR with primers amplifying the transgenes mag or αLNNd. Correct targeting was further confirmed with a PCR using a primer recognizing a genomic sequence outside of the upstream homology region of the targeting vector and a second primer in mag or αLNNd. Targeted founder mice were crossed to homozygosity and loss of wild-type *Rosa26* allele was confirmed by PCR. Genotyping was performed with primers on the *Rosa26* wild-type allele: 5’-AAG GGA GCT GCA GTG GAG TA and 5’-CCG AAA ATC TGT GGG AAG TC; on aLNNd: 5’-AGC TGA TCC GGA ACC CTT AA and 5’-GGA TGG CGC TCT CTA GGA TT; on mag: 5’-AAG GGA GCT GCA GTG GAG TA and 5’-CCG AAA ATC TGT GGG AAG TC.

To express mag and αLNNd ubiquitously, mice were crossed with CAG-Cre-ER™ mice (Hayashi & McMahon, 2002) [B6.Cg-Tg(CAG-cre/Esr1*)5Amc/J available from the Jackson Laboratory]. CAG-Cre-ER™ mice were crossed with tdTomato-expressing reporter mice [Ai9; B6;129S6-Gt(ROSA)26Sortm9(CAG-tdTomato)Hze/J (Madisen *et al*., 2010) available from the Jackson Laboratory] to control for Cre-mediated recombination in skeletal muscle. For tamoxifen application, tamoxifen (Sigma; T5648-1G) was dissolved in corn oil (Sigma; C8267) at a concentration of 20 mg/ml by shaking overnight at 37°C and administrated at a dose of 75 mg/kg by i.p. injection once every 24 h for a total of 5 consecutive days. 14 days after the first injection, tissue was collected and recombination and removal of the LSL cassette was confirmed on purified DNA using primers flanking the LSL cassette: 5’-GCT GGT TAT TGT GCT GTC TCA TC and 5’-TGC ACT TAA CGC GTA CAA GG.

All mice analyzed were from breedings of mice heterozygous for the knockout allele in the *Lama2* locus, hemizygous for CAG-Cre-ER™ and homozygous for LSL-αLNNd or LSL-mag alleles, respectively. This strategy allowed to receive all genotypes from the same breeding and use littermates as controls. Control mice were always Cre-negative and heterozygous for LSL-αLNNd and LSL-mag; for the *Lama2* locus, controls were either wild-type or heterozygous. Unless otherwise indicated, female and male mice were used. To ensure optimal access of the dystrophic mice to water and food, all cages were supplied with long-necked water bottles and wet food from weaning onwards. All mouse experiments were performed according to the federal guidelines for animal experimentation and approved by the authorities of the Canton of Basel-Stadt.

### Antibodies

For immunostaining and Western blot analysis, the following antibodies were used: α-actinin (Sigma, catalog no. A7732; 1:5000), agrin (for detection of mag) (produced in-house (Eusebio *et al*, 2003); 1:5,000 for Western blots, 1:200 for immunostainings) or (R&D System catalog no. AF550; 5 μg/ml for immunostainings), laminin-α1 (for detection of αLNNd by Western blot) R&D Systems, catalog no. AF4187; 1:2,000), αLNNd (for detection of αLNNd by immunostainings, previously described (McKee *et al*., 2017); 1:100), laminin-α2 (Sigma, catalog no. L0663, clone 4H8-2; 1:500), laminin-α4 (previously described (McKee *et al*., 2017); 1:1,000 for Western blots, 1:200 for immunostainings), laminin-β1γ1 (Sigma, catalog no. L9393; 1:100), GAPDH (Cell Signaling, catalog no. 2118, 1:1,000).

### Immunostainings

Fresh frozen muscle or sciatic nerve tissue was cryo-sectioned (10 μm) and fixed with 4% paraformaldehyde in PBS (4% PFA/PBS) or stained non-fixed. Sections were incubated in blocking solutions (5% donkey serum and 0.3% triton-X in PBS) for 1 h at room temperature and then incubated overnight at 4°C with primary antibodies, diluted in 5% donkey serum in PBS. Sections were washed three times with PBS and incubated at room temperature for 1 h with the appropriate secondary antibodies. Images were acquired with an Olympus iX81 microscope, a Zeiss Axio Scan.Z1 Slide Scanner or a Zeiss point scanning Confocal microscope LSM700.

### Protein extraction and Western blot analysis

For total muscle extracts, frozen muscles were pulverized in liquid nitrogen, lysed in modified RIPA buffer (50 mM Tris-HCl pH 8.0, 150 mM NaCl, 1% NP-40, 0.5% deoxycholate, 0.1% SDS, 20 mM EDTA; protease inhibitors), sonicated and incubated for 2 h at 4°C with head-over-head rotation. Insoluble material was removed by centrifugation (16,000g, 30 min, 4°C). After adjustment of protein concentrations (determined by BCA assay), Laemmli buffer was added, samples incubated for 5 min at 95°C and subjected to standard Western blot procedures.

### Histology and histological quantifications of muscle

Muscles were mounted on 7% gum tragacanth (Sigma), and rapidly frozen in 2-methylbutane that was cooled in liquid nitrogen (−150°C). Cross-sections of 10 μm thickness were cut on a cryostat. General histology was assessed after tissue fixation with 4% paraformaldehyde by H&E staining (Merck) or Picro Sirius Red stain (Direct Red 80 (Sigma) in picric acid solution). Images were acquired with an Olympus iX81 microscope using cellSens software (Olympus). The muscle fiber diameter was quantified using the minimum distance of parallel tangents at opposing particle borders (minimal “Feret’s diameter”), as described previously (Briguet *et al*, 2004). For fiber number and fiber size analysis, complete mid-belly cross sections were evaluated using an in-house-customized, Fiji-based version of Myosoft (Encarnacion-Rivera *et al*, 2020; Schleicher, 2022). The algorithms used are available at <u>10.5281/zenodo.6469872</u>.

### Histology and morphometric analysis of sciatic nerves

Semithin and ultrathin morphological experiments were performed as described previously (Porrello *et al*, 2014). In brief, sciatic nerves were dissected and fixed with 2.5% glutaraldehyde in 0.12 M phosphate buffer pH7.4, post-fixed with 1% osmium tetroxide, and embedded in epon (Sigma catalog no. 45359). Semithin sections (1 μm) were stained with 0.1% toluidine blue and examined by light microscopy on a BX51 Olympus microscope. To perform morphometric analysis, images of cross sections were obtained from corresponding levels of the sciatic nerve by a digital camera (DCF7000T, Leica) with 100x objective. Five images per animal were acquired and analyzed using the Leica QWin Software (Leica Microsystem, Milan). The G-ratio (axon diameter/fiber diameter) was determined by dividing the mean diameter of an axon by the mean diameter of the same fiber (axon plus myelin). At least 350 randomly chosen fibers per animal were examined. Ultrathin sections (70-80 nm thick) were stained with uranyl acetate and lead citrate and examined by electron microscopy (FEI Talos L120C G2 Transmission Electron Microscope). At least 20 images per animal, randomly photographed, were acquired and analyzed using ImageJ (National Institutes of Health) software.

### Cell culture

COS7 cells were cultured in DMEM medium (Invitrogen) in the presence of 10% fetal bovine serum (Gibco). Transfections of LSL-mag, LSL-αLNNd and CAG-Cre (Matsuda & Cepko, 2007) (Addgene Cat. 13775) plasmids were performed with Lipofectamin 2000 (Invitrogen) following the manufacturer’s instructions. For the analysis of proteins in cell-conditioned medium, medium was replaced 24 h after transfection with serum-free DMEM. 48 h after transfection, conditioned medium was collected and COS7 cells were lysed. Both samples were subjected to Western blot analysis.

### Grip strength measurement

Forelimb grip strength was assessed according SOP MDC1A_M.2.2.001 (TREAT-NMD) using a grip strength meter (Columbus) equipped with a trapeze bar. Mice were lifted on the tail towards the bar until the mouse gripped it with both forepaws and then gently moved away at constant speed until its grip was lost. The peak force was calculated as the mean of the three best trials out of six consecutive pulls.

### Gait analysis

Gait analysis was performed using the CatWalk XT system (Noldus) following the manufacturer’s instructions. Mice were placed in an enclosed illuminated walkway on a glass plate, allowed to move freely in both directions and their footprints recorded by a high-speed video camera positioned underneath the walkway. Runs were classified as compliant when mice crossed a defined 40 cm distance in the walkway within 10 s and a maximum speed variation of 60%. Three compliant runs were recorded for each mouse and averaged for statistical analysis. For *dy*^*W*^/*dy*^*W*^ mice, only three out of six mice measured were able to perform compliant runs, whereas compliant runs were obtained for all assessed control and *dy*^*W*^/*dy*^*W*^ CAG-DL mice.

### Assessment of motor coordination (rotarod assay)

Motor coordination of mice was assessed using a rotarod device (Ugo Basile). Mice were trained for 2 consecutive days by placing them 3 times for 1 min on the rotarod rotating at a constant speed of 5 rpm. On the third consecutive day, performance was assessed on accelerating rod (5-40 rpm within 5 min) for a maximal duration of 400 s. The test was performed 3 times with minimal rest period of 10 min between trials. The time on the rod (latency to fall) was averaged for the three trials.

### In vitro muscle force measurement

In vitro muscle force measurements were performed on isolated EDL muscle using the 1200A Isolated Muscle System (Aurora Scientific) in an organ bath at 30°C containing Ringer solution (137 mM NaCl, 24 mM NaHCO_3_, 11 mM glucose, 5 mM KCl, 2 mM CaCl_2_, 1 mM MgSO_4_, and 1 mM NaH_2_PO_4_) constantly oxygenated with 95% O_2_/5% CO_2_. After muscles were adjusted to the optimum muscle length (Lo), muscles were stimulated with electrical pulse at 15 V and achieved peak twitch force (Pt) recorded. Peak tetanic force (Po) was assessed as maximal force with 500 ms stimulation at 10-250 Hz. Specific twitch and tetanic forces were calculated by normalization to the cross-sectional area (CSA) by using the formula CSA (mm2) = muscle wet weight (mg)/[fiber length (lf, mm) × 1.06 mg/mm3], with lf = lo × 0.44 for EDL as described (Brooks & Faulkner, 1988).

### Study design and Statistical analysis

Mice were randomly allocated to experimental groups. Evaluations of immunohistochemistry, muscle histology, muscle function and behavior assays were performed by investigators blinded to the specific sample. Statistical analysis was performed using unpaired, two-tailed Student’s t test for comparisons of two groups. For the comparisons of more than two groups, one-way ANOVA followed by Bonferroni post-hoc test was used or, if unequal variance between groups was detected, one-way ANOVA with Dunnett’s multiple comparisons test. We assumed normal distribution of the variables analyzed. All statistical tests were performed using Prism version 9 (GraphPad Software).

## Supporting information

Movie S1. 14-week-old dyW/dyW, dyW/dyW CAG-DL and control mouse

Movie S2. 4-week-old dy3K/dy3K, dy3K/dy3K CAG-DL and control mouse

Movie S3. 2-month-old and 16-month-old dy3K/dy3K CAG-DL mouse

## ACKNOWLEDGMENTS

We thank Dr. Kai D. Schleicher from the Imaging Core Facility for customizing Myosoft, Heide Oller from the Center for Transgenic Models for help with transgenic mice generation and Paola Podini for help with EM images. The work in the laboratory of MAR was supported by the Cantons of Basel-Stadt and Basel-Landschaft, by Cure CMD, the Swiss Foundation for Research on Muscle Disease (FSRMM) and by Innosuisse (grant no. 34870.1 IP-LS). Work in the laboratory of SCP was supported by the European Joint Programme on Rare Diseases (MYOCITY).

## AUTHOR CONTRIBUTIONS

JRR, SCP and MAR designed experiments. JRR, EP and SL performed experiments and analyzed the data. JRR and MAR wrote the manuscript and all the co-authors commented.

## REFERENCES

Bentzinger CF, Barzaghi P, Lin S, Ruegg MA (2005) Overexpression of mini-agrin in skeletal muscle increases muscle integrity and regenerative capacity in laminin-alpha2-deficient mice. FASEB J 19: 934–942

Briguet A, Courdier-Fruh I, Foster M, Meier T, Magyar JP (2004) Histological parameters for the quantitative assessment of muscular dystrophy in the mdx-mouse. Neuromuscul Disord 14: 675–682

Brooks SV, Faulkner JA (1988) Contractile properties of skeletal muscles from young, adult and aged mice. J Physiol 404: 71–82

Chu VT, Weber T, Graf R, Sommermann T, Petsch K, Sack U, Volchkov P, Rajewsky K, Kuhn R (2016) Efficient generation of Rosa26 knock-in mice using CRISPR/Cas9 in C57BL/6 zygotes. BMC Biotechnol 16: 4

Domogatskaya A, Rodin S, Tryggvason K (2012) Functional diversity of laminins. Annu Rev Cell Dev Biol 28: 523–553

Encarnacion-Rivera L, Foltz S, Hartzell HC, Choo H (2020) Myosoft: An automated muscle histology analysis tool using machine learning algorithm utilizing FIJI/ImageJ software. PLoS One 15: e0229041

Eusebio A, Oliveri F, Barzaghi P, Ruegg MA (2003) Expression of mouse agrin in normal, denervated and dystrophic muscle. Neuromuscul Disord 13: 408–415

Feltri ML, Poitelon Y, Previtali SC (2016) How Schwann Cells Sort Axons: New Concepts. Neuroscientist 22: 252–265

Gawlik K, Miyagoe-Suzuki Y, Ekblom P, Takeda S, Durbeej M (2004) Laminin alpha1 chain reduces muscular dystrophy in laminin alpha2 chain deficient mice. Hum Mol Genet 13: 1775–1784

Gawlik KI, Durbeej M (2020) A Family of Laminin alpha2 Chain-Deficient Mouse Mutants: Advancing the Research on LAMA2-CMD. Front Mol Neurosci 13: 59

Haueter S, Kawasumi M, Asner I, Brykczynska U, Cinelli P, Moisyadi S, Burki K, Peters AH, Pelczar P (2010) Genetic vasectomy-overexpression of Prm1-EGFP fusion protein in elongating spermatids causes dominant male sterility in mice. Genesis 48: 151–160

Hayashi S, McMahon AP (2002) Efficient recombination in diverse tissues by a tamoxifen-inducible form of Cre: a tool for temporally regulated gene activation/inactivation in the mouse. Dev Biol 244: 305–318

Hoy SM (2019) Onasemnogene Abeparvovec: First Global Approval. Drugs 79: 1255–1262

Kong L, Valdivia DO, Simon CM, Hassinan CW, Delestree N, Ramos DM, Park JH, Pilato CM, Xu X, Crowder M et al (2021) Impaired prenatal motor axon development necessitates early therapeutic intervention in severe SMA. Sci Transl Med 13

Kuang W, Xu H, Vachon PH, Engvall E (1998a) Disruption of the lama2 gene in embryonic stem cells: laminin alpha 2 is necessary for sustenance of mature muscle cells. Exp Cell Res 241: 117–125

Kuang W, Xu H, Vachon PH, Liu L, Loechel F, Wewer UM, Engvall E (1998b) Merosin-deficient congenital muscular dystrophy. Partial genetic correction in two mouse models. J Clin Invest 102: 844–852

Madisen L, Zwingman TA, Sunkin SM, Oh SW, Zariwala HA, Gu H, Ng LL, Palmiter RD, Hawrylycz MJ, Jones AR et al (2010) A robust and high-throughput Cre reporting and characterization system for the whole mouse brain. Nat Neurosci 13: 133–140

Manini A, Abati E, Nuredini A, Corti S, Comi GP (2021) Adeno-Associated Virus (AAV)-Mediated Gene Therapy for Duchenne Muscular Dystrophy: The Issue of Transgene Persistence. Front Neurol 12: 814174

Matsuda T, Cepko CL (2007) Controlled expression of transgenes introduced by in vivo electroporation. Proc Natl Acad Sci U S A 104: 1027–1032

McKee KK, Crosson SC, Meinen S, Reinhard JR, Ruegg MA, Yurchenco PD (2017) Chimeric protein repair of laminin polymerization ameliorates muscular dystrophy phenotype. J Clin Invest 127: 1075–1089

McKee KK, Harrison D, Capizzi S, Yurchenco PD (2007) Role of laminin terminal globular domains in basement membrane assembly. J Biol Chem 282: 21437–21447

Meinen S, Barzaghi P, Lin S, Lochmuller H, Ruegg MA (2007) Linker molecules between laminins and dystroglycan ameliorate laminin-alpha2-deficient muscular dystrophy at all disease stages. J Cell Biol 176: 979–993

Miyagoe Y, Hanaoka K, Nonaka I, Hayasaka M, Nabeshima Y, Arahata K, Nabeshima Y, Takeda S (1997) Laminin alpha2 chain-null mutant mice by targeted disruption of the Lama2 gene: a new model of merosin (laminin 2)-deficient congenital muscular dystrophy. FEBS Lett 415: 33–39

Moll J, Barzaghi P, Lin S, Bezakova G, Lochmuller H, Engvall E, Muller U, Ruegg MA (2001) An agrin minigene rescues dystrophic symptoms in a mouse model for congenital muscular dystrophy. Nature 413: 302–307

Nguyen Q, Lim KRQ, Yokota T (2019) Current understanding and treatment of cardiac and skeletal muscle pathology in laminin-alpha2 chain-deficient congenital muscular dystrophy. Appl Clin Genet 12: 113–130

Patton BL, Connoll AM, Martin PT, Cunningham JM, Mehta S, Pestronk A, Miner JH, Sanes JR (1999) Distribution of ten laminin chains in dystrophic and regenerating muscles. Neuromuscul Disord 9: 423–433

Patton BL, Wang B, Tarumi YS, Seburn KL, Burgess RW (2008) A single point mutation in the LN domain of LAMA2 causes muscular dystrophy and peripheral amyelination. J Cell Sci 121: 1593–1604

Porrello E, Rivellini C, Dina G, Triolo D, Del Carro U, Ungaro D, Panattoni M, Feltri ML, Wrabetz L, Pardi R et al (2014) Jab1 regulates Schwann cell proliferation and axonal sorting through p27. J Exp Med 211: 29–43

Previtali SC, Zambon AA (2020) LAMA2 Neuropathies: Human Findings and Pathomechanisms From Mouse Models. Front Mol Neurosci 13: 60

Proske U, Gandevia SC (2012) The proprioceptive senses: their roles in signaling body shape, body position and movement, and muscle force. Physiol Rev 92: 1651–1697

Qiao C, Dai Y, Nikolova VD, Jin Q, Li J, Xiao B, Li J, Moy SS, Xiao X (2018) Amelioration of Muscle and Nerve Pathology in LAMA2 Muscular Dystrophy by AAV9-Mini-Agrin. Mol Ther Methods Clin Dev 9: 47–56

Reinhard JR, Lin S, McKee KK, Meinen S, Crosson SC, Sury M, Hobbs S, Maier G, Yurchenco PD, Ruegg MA (2017) Linker proteins restore basement membrane and correct LAMA2-related muscular dystrophy in mice. Sci Transl Med 9

Sarkozy A, Foley AR, Zambon AA, Bonnemann CG, Muntoni F (2020) LAMA2-Related Dystrophies: Clinical Phenotypes, Disease Biomarkers, and Clinical Trial Readiness. Front Mol Neurosci 13: 123

Sasaki T, Giltay R, Talts U, Timpl R, Talts JF (2002) Expression and distribution of laminin alpha1 and alpha2 chains in embryonic and adult mouse tissues: an immunochemical approach. Exp Cell Res 275: 185–199

Schleicher KD (2022) imcf/myosoft-imcf: myosoft-imcf-1.0.0. Zenodo: 10.5281/zenodo.6469873

Xu H, Wu XR, Wewer UM, Engvall E (1994) Murine muscular dystrophy caused by a mutation in the laminin alpha 2 (Lama2) gene. Nat Genet 8: 297–302

Yao X, Zhang M, Wang X, Ying W, Hu X, Dai P, Meng F, Shi L, Sun Y, Yao N et al (2018) Tild-CRISPR Allows for Efficient and Precise Gene Knockin in Mouse and Human Cells. Dev Cell 45: 526–536 e525

Yurchenco PD (2011) Basement membranes: cell scaffoldings and signaling platforms. Cold Spring Harb Perspect Biol 3

Yurchenco PD, McKee KK, Reinhard JR, Ruegg MA (2018) Laminin-deficient muscular dystrophy: Molecular pathogenesis and structural repair strategies. Matrix Biol 71-72: 174–187

