## Supplementary material for "Nerve pathology is prevented by linker proteins in mouse models for *LAMA2*-related muscular dystrophy"

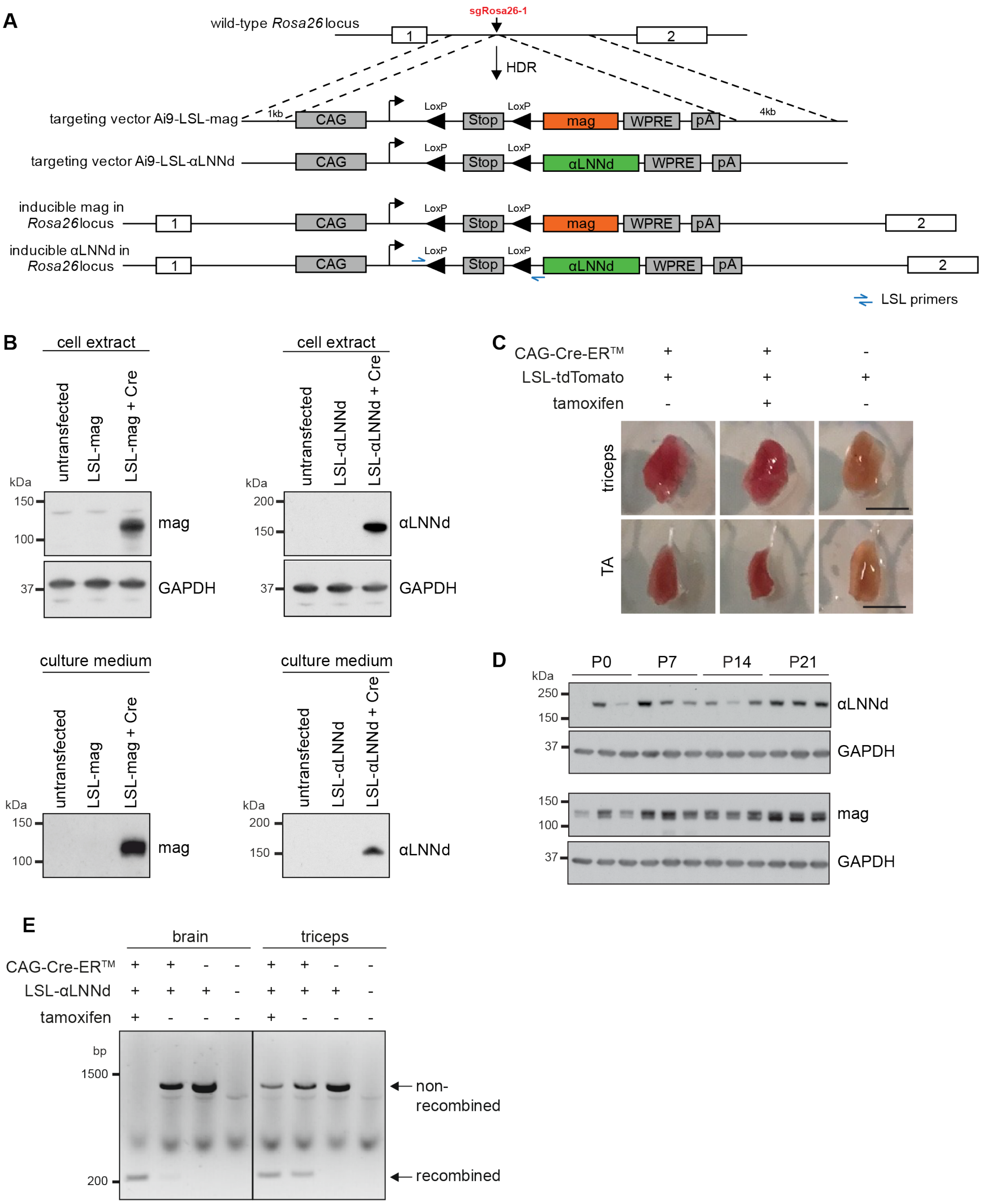


**Figure S1. Generation and characterization of mice expressing the linker proteins ubiquitously (CAG-DL mice).**

**A** Schematic of the Cre-dependent LSL-mag and LSL-αLNNd targeting constructs and the position of the sgRosa26-1 (Chu *et al.*, 2016) used for cutting the *Rosa26* locus.

**B** Western blot analysis of cell extracts and cell culture medium from COS7 cells transfected with indicated plasmids. GAPDH was used as loading control.

**C** Expression of tdTomato in *triceps brachii* (TRC) and *tibialis anterior* (TA) muscle of 5-week-old mice with the indicated genotype and upon injection of tamoxifen. Note that tdTomato is expressed in both muscles irrespective of tamoxifen treatment.

**D** Western blot analysis of lysate from TRC muscle of CAG-DL mice not injected with tamoxifen from postnatal day zero (P0) to P21. GAPDH was used as loading control.

**E** PCR on genomic DNA purified from brain or *triceps brachii* (triceps) muscle from 8-week-old mice using the indicated primers in (**A**). PCR products before (non-recombined) and after Cre-mediated recombination (recombined) of the LSL cassette are indicated. Note the high level of recombination observed in triceps muscle in CAG-CreER$^{TM}$-positive mice that were not treated with tamoxifen.

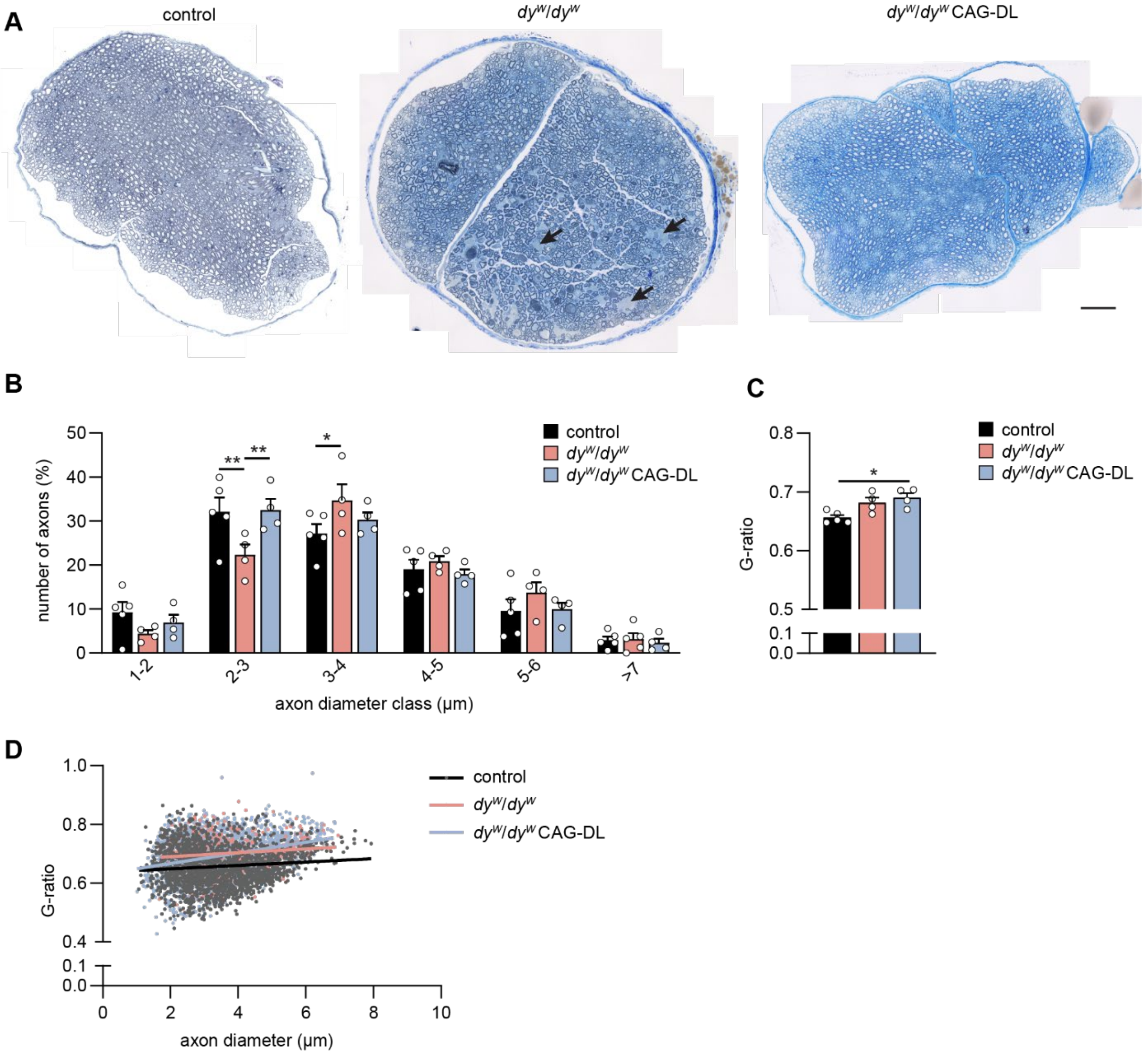


**Figure S2. Quantification of nerve pathology.**

**A** Reconstruction of the entire nerve using toluidine blue-stained, semi-thin sections of the sciatic nerve from 8-week-old mice of the indicated genotype. Compared to controls, *dy*$^{W}$/*dy*$^{W}$ mice contain large regions/bundles of non-myelinated axons (arrows). Scale bar: 40 µm

**B-D** Quantification of myelination in sciatic nerves from 8-week-old mice of the indicated genotype. Some changes between the genotypes are seen in the axon diameters (**B**), the G-ratio (**C**) and the scatter plot of G-ratio values versus axon diameter (**D**).

Data are mean ± SEM. **P* < 0.05; ***P* < 0.01; ****P* < 0.001; by one-way ANOVA with Bonferroni post hoc test (**B-C**). Lines indicate the linear regression (**D**). N = 4-5 mice per group.

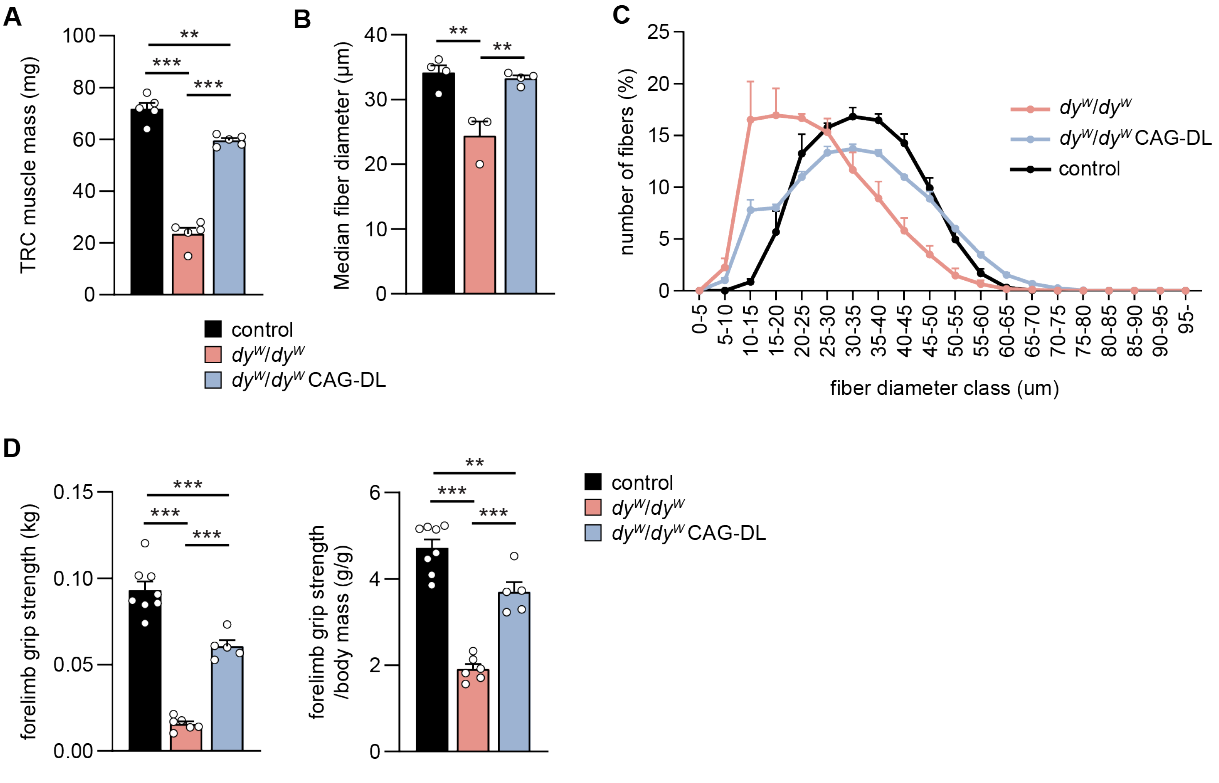


**Figure S3. Quantification of muscle histology of triceps muscle and forelimb grip strength.**

**A-C** Quantification of the *triceps brachii* (TRC) muscle mass (**A**) and its muscle fiber diameters (**B, C**) from 8-week-old female mice of the indicated genotype.

**D** Grip strength performance of 8-week-old female mice of the indicated genotype.

Data are mean ± SEM. *$P < 0.05$; *$P < 0.005$; ***$P < 0.001$; by one-way ANOVA with Bonferroni post hoc test. N = 3-8 mice per group.

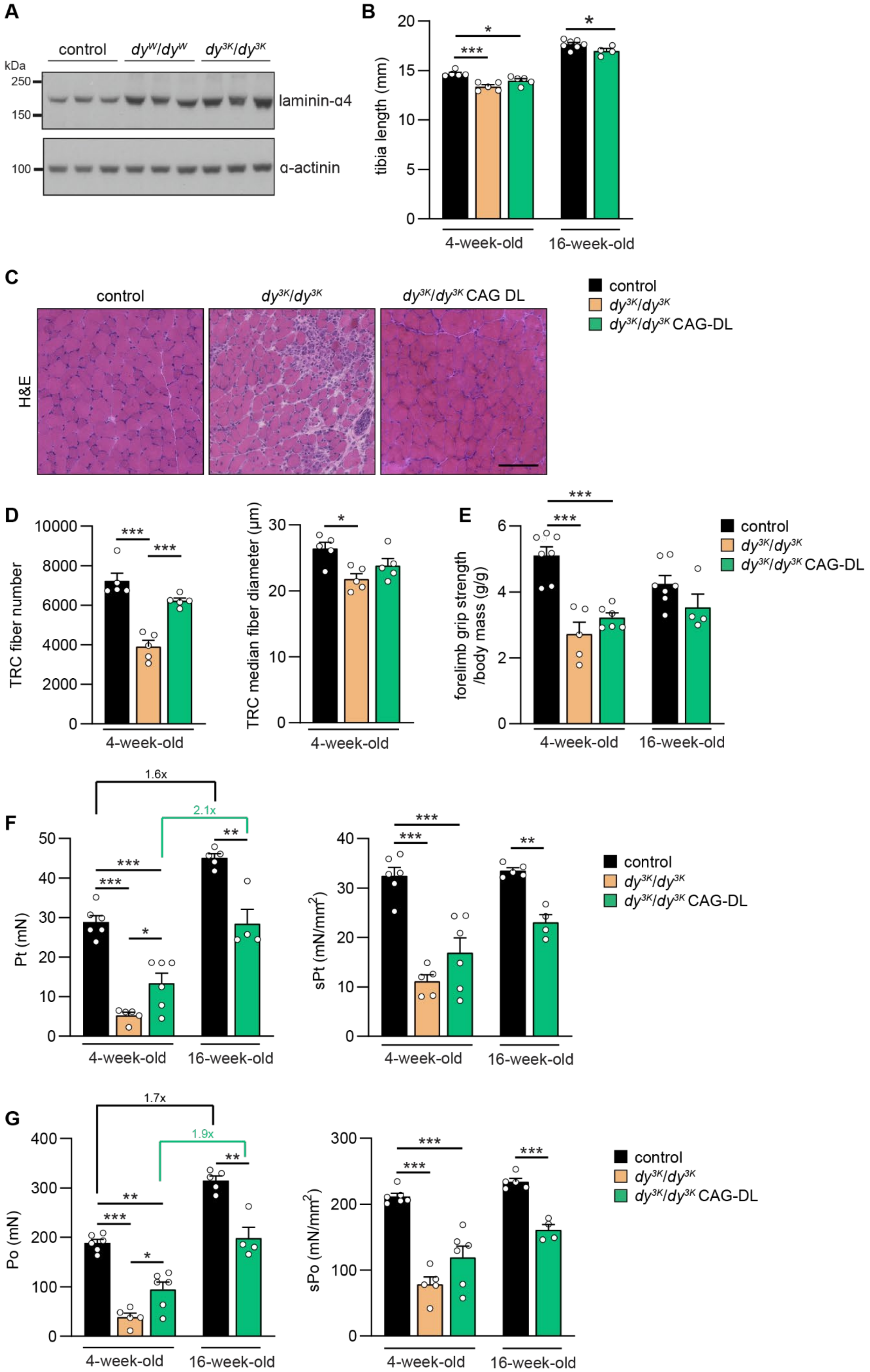
A
control
dyW/dyW
dy3K/dy3K
kDa
250
150
100
laminin-α4
α-actinin
B
tibia length (mm)
4-week-old
16-week-old
C
control
dy3K/dy3K
dy3K/dy3K CAG DL
H&E
control
dy3K/dy3K
dy3K/dy3K CAG-DL
D
TRC fiber number
TRC median fiber diameter (μm)
4-week-old
E
forelimb grip strength /body mass (g/g)
4-week-old
16-week-old
F
1.6x
2.1x
Pt (mN)
sPt (mN/mm2)
4-week-old
16-week-old
G
1.7x
1.9x
Po (mN)
sPo (mN/mm2)
4-week-old
16-week-old

**Figure S4. Treatment effect in *dy^3K^/dy^3K^* mice.**

**A** Western blot analysis for laminin-α4 in 4-week-old mice of the indicated genotype. α-actinin was used as loading control.

**B** Tibia length of 4- and 16-week-old mice of the indicated genotype.

**C** Hematoxylin and Eosin (H&E) of 4-week-old *tibialis anterior* (TA) muscle of the indicated genotype.

**D** Quantification of *triceps brachii* (TRC) muscle fiber number and diameter of 4-week-old female mice.

**E** Grip strength performance normalized to body weight of 4- and 16-week-old female mice of the indicated genotype.

**F**, **G** Twitch (**F**) or tetanic force (**G**) of *extensor digitorum longus* (EDL) muscle from 4- and 16-week-old female mice of the indicated genotype. Shown are absolute (left) and specific (right) force.

Data are mean ± SEM. *$P$ < 0.05; *$P$ < 0.005; ***$P$ < 0.001; by one-way ANOVA with Bonferroni post hoc test. N = 4-6 mice per group.

**Movie S1.** 14-week-old $dy^{W}/dy^{W}$, $dy^{W}/dy^{W}$ CAG-DL and control mouse.

**Movie S2.** 4-week-old $dy^{3K}/dy^{3K}$, $dy^{3K}/dy^{3K}$ CAG-DL and control mouse.

**Movie S3.** 2-month-old and 16-month-old $dy^{3K}/dy^{3K}$ CAG-DL mouse.
